# Community health and human-animal contacts on the edges of Bwindi Impenetrable National Park, Uganda

**DOI:** 10.1101/2021.07.23.453553

**Authors:** Renata L. Muylaert, Ben Davidson, Alex Ngabirano, Gladys Kalema-Zikusoka, Hayley MacGregor, James O. Lloyd-Smith, Ahmed Fayaz, Matthew A. Knox, David T. S. Hayman

**Author notes:** These authors contributed equally to this work (Renata L. Muylaert, Ben Davidson, David T. S. Hayman).

## Abstract

Cross-species transmission of pathogens is intimately linked to human and environmental health. With limited healthcare and challenging living conditions, people living in poverty may be particularly susceptible to endemic and emerging diseases. Similarly, wildlife is impacted by human influences, including pathogen sharing, especially for species in close contact with people and domesticated animals. Here we investigate human and animal contacts and human health in a community living around the Bwindi Impenetrable National Park (BINP), Uganda. We used contact and health survey data to identify opportunities for cross-species pathogen transmission, focusing mostly on people and the endangered mountain gorilla. We conducted a survey with background questions and self-reported diaries to investigate 100 participants’ health, such as symptoms and behaviours, and contact patterns, including direct contacts and sightings over a week. Contacts were revealed through networks, including humans, domestic, peri-domestic, and wild animals for 1) network of contacts seen in the week of background questionnaire completion, 2) network of contacts seen during the diary week. Participants frequently felt unwell during the study, reporting from one to 10 disease symptoms at different intensity levels (maximum of seven symptoms in one day), with severe symptoms comprising 6.4% of the diary records and tiredness and headaches the most common symptoms. Besides human-human contacts, direct contacts with livestock and peri-domestic animals were the most common. Wildlife contacts were the rarest, including one direct contact with gorilla with a concerning timeline of reported symptoms. The contact networks were moderately connected and revealing a preference in contacts within the same species or taxon and within their groups. Despite sightings of wildlife being much more common than touching, one participant declared direct contact with a mountain gorilla during the week. Gorillas were seen very close to six animal taxa (including themselves) considering all interaction types, mostly seen closer to other gorillas, but also people and domestic animals. Our findings reveal a local human population with recurrent symptoms of illness in a location with intense exposure to factors that can increase pathogen transmission, such as direct contact with domestic and wild animals and proximity among animal species. Despite significant biases and study limitations, the information generated here can guide future studies, such as models for disease spread and One Health interventions.

## Introduction

The COVID-19 pandemic, now the ‘poster child’ of the impact of an emerging infectious disease, is suspected to have originated from a virus circulating in wild mammals [1]. Ebola virus disease, HIV/AIDS, and SARS are all infamous emerging infectious diseases with origins in wildlife. In fact, most human infections have their origins in wild or domestic animals. Therefore, despite providing many benefits to health—such as overall well-being and services, including medicinal herbs and provision of other natural resources [2–4]—green natural areas can also pose risks to human health, especially when living in proximity to green areas is linked to social inequality and low-socio-economic status.

In low to middle-income countries, and particularly in tropical regions, people also still suffer from a high burden of numerous and treatable endemic diseases. These infections may be human-adapted (e.g., malaria [5]) or zoonotic (e.g. leptospirosis [6]). Contacts between humans and other animals can, therefore, pose hazards to human health, but the converse is also true [7]. Human behaviour has contributed to species extinction rates much higher than pre-Anthropocene levels [8]. Habitat destruction, climate change, pollution, species invasion, wildlife trade, and overharvesting [9–11] can negatively impact ecosystems, leading to the declining health of many species [12]. In some cases, this may be in the form of increased risk of pathogen transmission, with many examples where infectious diseases have contributed to the decline of endangered wildlife species [13,14].

Disease risk is especially important in non-human great apes, such as the endangered chimpanzees, bonobos (*Pan paniscus*), the Western lowland gorilla (*Gorilla gorilla gorilla*), Grauer’s gorilla (*Gorilla beringei graueri*), and the mountain gorilla (*Gorilla beringei beringei*). They are all endangered or critically endangered species [15] threatened by disease, poaching, and habitat loss. The plight of non-human great apes is even more critical because their close relatedness to humans may pose a higher risk for cross-species transmission of infections [16,17]. Human respiratory viruses can be transmitted to gorilla populations from people, as in the case of Human Respiratory Syncytial Virus [18].

Ebola virus disease (EVD) outbreaks in Central Africa have killed thousands of both people and apes, with uncertain transmission pathways. These outbreaks have decimated the critically endangered Eastern Lowland gorilla (*Gorilla beringei graueri*) population, reaching 90% mortality in some areas [19]. Analyses of human EVD outbreaks suggest that land-use changes, particularly forest fragmentation, may be increasing the risk of these outbreaks [20,21]. The human population density is high in many African regions that share an interface with non-human great apes’ habitat, such as in Bwindi Impenetrable National Park (BINP) and the Virunga Volcanoes. Habitat degradation and human encroachment—the land bordering BINP has an average population density of 300-400 people/km^2^ [22], much higher than the average for Uganda (229 people/km^2^, [23])—and ecotourism increase human-wildlife contact.

Communities around BINP have had to deal with civil unrest, including after the displacement of human populations by the creation of the park [24]. Precarious livelihoods are common around the park. Yet, the area is an ecotourism destination with frequent visits to habituated groups of apes [25]. These factors, along with high inbreeding levels [26], probably increase the frequency and severity of disease outbreaks in mountain gorillas [27,28] due to the increased susceptibility to and transmission of infections via increasing contact rates.

More than 20 disease-causing pathogens are transmitted from humans to great apes [25]. These pathogens include several viruses [18,29,30], bacteria, ectoparasites and endoparasites [31], leading to diseases such as giardiasis and cryptosporidiosis [32]. Close contact, within a few meters of a coughing or sneezing individual, would be enough for pathogen transmission from one species to another. Moreover, pathogens spread by the faecal-oral route can be readily shared among species that overlap spatially and share resources such as waterways. On the other hand, regarding the gorilla-human interface, there are no reports of HIV transmission between humans and gorillas. However, HIV causes immunosuppression in humans, leaving people susceptible to tuberculosis and other respiratory diseases, which pose a threat to gorilla populations [25,33,34]. Nevertheless, there are immunodeficiency viruses that had their ancestry in apes [35], such as SIVcpz from chimpanzees in Cameroon [36], which is thought to be the ancestral group for the pandemic HIV-1 group M. Thus, it is essential to investigate contact patterns between humans and other primates at the boundaries of natural areas.

Interdisciplinary frameworks consider key ultimate and proximate components that drive cross-species infection transmission to understand the process of disease emergence [37,38]. Studying and addressing these components requires multidisciplinary teams comprising local community members, ecologists, anthropologists, physicians, and more [39,40]. Ultimate drivers include broad environmental, social and politico-economic drivers operating at different timescales and spatial scales [38,41]. Living conditions and higher-level structural drivers, beyond the actions of individuals, may have a large impact on people’s livelihoods and the outcomes of the interactions between humans and nature. Human communities in poor regions in proximity to natural areas frequently suffer from various diseases and take the toll of inadequate health facilities, the precarity of livelihoods, food insecurity, illiteracy, and concurrent proximity to livestock and wildlife, making them susceptible to cross-species transmission. The proximate drivers comprise the 1) prevalence in the reservoir, which falls in the scope of disease ecology; 2) contact rates, a prevalent knowledge gap for many systems; and 3) probability of infection given exposure, which falls in the domain of microbial risk assessment.

Regarding contact rates, proximate driver 2, quantifying behaviour and contact patterns to inform disease spread models can be challenging. Large projects attempting to measure demographics and patterns of human-human contact have been conducted worldwide at small [42] and large scales [43], with essential applications in epidemiological models [44]. While identifying human contact patterns is important to predict onward disease spread, contact with wildlife and livestock is a crucial factor allowing pathogen cross-species transmission to happen. However, data sets describing human-animal contacts together with animal-animal contacts (such as Narat et al., in Central Africa [42]) are rarely reported [45,46]. Contact data between numerous species can be cumbersome to gather by direct observation but can be estimated from survey participants’ perspectives, whereas self-declared information can reveal relevant general contact patterns and health. Questionnaire research is hardly ever objective [47]; nonetheless, self-declared information can be used as feedback to guide actions most needed in small communities. Interventions like keeping distance from wildlife and handwashing before preparing meals can greatly reduce the cross-over of disease-causing pathogens [48,49]. Initiatives to promote these types of needed actions have been made in the BINP, which economically relies heavily on subsistence agriculture [50]. In this ecosystem, close relationships people have with other people, their livestock and other animals can be explored through social science approaches and measured by further focused enquiry informed by conceptual models of cross-species transmission [37,39,40]. Understanding the potential contacts between humans and other animals and their network structure can help inform the control measures required to manage and prevent future disease outbreaks.

This study used questionnaires to investigate people’s health and potential risk factors for infectious diseases investigating the contacts among humans, domestic, peri-domestic and wildlife, with a particular focus on humans and the endangered mountain gorilla. We describe contact patterns in humans and other animals living close to BINP through self-reported surveys for one week. We analysed contact patterns for 1) human-human contacts, including the participants’ social contacts and their reported observations; 2) animal-human contacts, including direct contacts and sightings and 3) their observation of animal-animal contacts in proximity to the animal-human interface. We expected human contact patterns to follow patterns reported previously for southwestern Uganda [51]. We expected wildlife sightings would be much more common than touching, with the existence of close contact preferences within-taxon and within groups (domestic, peri-domestic, people, and wild animals). Our questionnaire provides insights on community health and human-animal interface in two villages near BINP.

## Methods

### Location and context

This study took place in Mukono parish at Bwindi Impenetrable National Park’s Buhoma sector in Uganda, between June and August 2018. Questionnaires were completed by local villagers living next to the northwest border of BINP, Uganda. BINP covers an area of 331 km^2^ in southwestern Uganda (Fig 1A, Fig 1B), and the villages are adjacent to BINP at the north-western edge of the park. This community was selected because it represents a scenario with risks of infection for both humans and wildlife where One Health initiatives have been put in practice [34]. People in the community generally have very low incomes, poor public health, and live in farmland areas adjacent to the highly biodiverse forests (Fig 1A) [34,52].

**Figure 1.** **A. Hard border between the BINP forests and agricultural land.** Photo credit: D. Hayman. **B. Study location around Bwindi National Park**, where questionnaires were applied to 100 people living in the Buhoma sector, Mukono Parish, Kanungu District, Southwestern Uganda. Source for shapefile: Mesa-IGAD geoportal http://mesa-geoportal.icpac.net/layers/geonode%3Abwindilulc2010 (accessed in July 2020).

### Health around BINP

In Buhoma, the human lifetime fertility rate is five births per woman for 2020, with a median population age of 15.7 [53]. About 42% of people live in poverty in 2020 and the literacy rate is 77% (males 83%, females 71%, data from 2018 for 15+ year-old people [53]). The disease burden in the local human community is high with over 10% mortality in under 5-year-old children [54]. The population around Bwindi has a high prevalence of infectious diseases where malaria, respiratory diseases and diarrhoea are very common [55]. Moreover, the majority of the diagnoses from the local hospital are human diseases that can also pose a threat to great apes [56,57]. Limited access to clean water and soap are associated with poor hygiene in the local community, which can favour the spread of diseases, such as COVID-19, typhoid and Hepatitis A [58]. Traditional healers practice in the community alongside the nearest hospital providing basic medical care, just 2.8 km from the Buhoma park headquarters, and a local store that sells antibiotics and other remedies off the shelf in the local village.

### Human-animal interface around BINP

BINP was a forest reserve until it was gazetted as National Park in 1991 [59,60]. The park is home to around 120 mammal species with ten species of primates, including the endangered mountain gorilla [61]. Other primates common to this area include chimpanzees, blue monkeys (*Cercopithecus mitis*, hereafter monkeys), Colobus monkeys (*Colobus* spp., hereafter Colobus monkeys) and olive baboons (*Papio anubis*, hereafter baboons). The formation of BINP led to local resentment due to inadequate community consultation and concern about loss of access to resources. Agreements and policy implementation plans (Integrated Conservation and Development) to ensure that the local communities would benefit from the forest were made, such as tourism revenue sharing [50]. The shared revenue is invested in infra-structure, health care, schools and livestock. Moreover, the creation of the Bwindi and Mgahinga Conservation Trust intends to cover part of community development expenses. However, there is still local concern regarding the distribution of the benefits to the community. The economic activities in the area centre on subsistence farming (agriculture, cattle, goats, pigs, and chicken [24,62]) and local living conditions are often inadequate (see S1 Fig and S2 Fig). The Bakiga—the predominant ethnic group living adjacent to Bwindi Impenetrable BINP forests in Buhoma sector [63]—are notably farmers, and traditionally the Batwa people lived in the interior of Bwindi Impenetrable forest as hunter-gatherers. Both Batwa and Bakiga were prevented from using BINP land in 1991; Bakiga could not open farming areas in BINP any longer and Batwa were forced to dwell outside the forest edge because of the restricted land use policies [24,59,60]. Batwa livelihoods now depend of exchanging labour for food, usually in other people’s lands [62].

Most people living around BINP are primarily subsistence farmers, rearing livestock and planting crops up to the forest edge (S2 Fig). This provides an active interface between the wildlife in the park, the local villagers and their livestock, and it is at this interface where pathogens can cross between humans, their domestic animals and wildlife [64]. With no buffer zone between the forest habitat of the national park and the surrounding area (Fig 1A), wildlife from the national park frequently enter the intensively cultivated farmland adjacent to the park to find food. A network of streams crosses through the forest and farmland, creating a common source of drinking water—another potential route of cross-contamination between people and animals. Moreover, despite the fact that hunting is illegal, there is evidence of poaching in the Park’s interior—for instance, 88 snares were destroyed in 2018 [65].

### Community organizations

Several organisations work locally aiding conservation and social development. Conservation Through Public Health (CTPH) is an organisation working at a community level and taking a conservation medicine approach at the human-animal-environment interface, with an overall focus on mountain gorilla conservation. CTPH works closely with local health providers (Bwindi Community Hospital) and the community, as well as with Uganda Wildlife Authority. CTPH promotes a One Health approach and supports the communities with guidance on family planning, hygiene, sanitation, infectious disease prevention and control, nutrition and sustainable agriculture. CTPH has a strong focus on education on the topics of forest conservation, risks of disease transmission among humans, gorillas and domestic animals, and monitoring homes visited by gorillas [52]. The Mountain Gorilla Veterinary Project, started by Diane Fossey in 1970, now the Gorilla Doctors, also runs a gorilla health program in the Virunga Volcanos and BINP [66]. Prior to the many collective efforts of these organisations and the application of active conservation strategies, the mountain gorilla population had dwindled to less than 250 animals. Now, decades later, the mountain gorilla population is over 1000 and the species was recently reclassified as endangered from critically endangered [15].

### Questionnaires

Questionnaires were designed by our multidisciplinary team, where the key information required included human-human, human-animal and animal-animal contacts. The questions were written in a manner appropriate to the local community and included pictures to provide clarity and to be as easy as possible to complete. Interviewees were recruited by AN, from CTPH, who is a local community member who has worked for CTPH since 2005. Small field trials were performed by DH and AN and the questions were refined and tested to ensure the data collected would be in a useable form and culturally acceptable. All further community engagement was by CTPH alone, led by AN. By asking the same question in different ways unreliable answers could be identified and these were excluded from the final analysis.

Participants were recruited through community meetings and visiting homes. Participant selection was through convenience sampling and based in part on participants’ willingness or perceived ability to complete the questionnaire. Only participants who gave their informed consent were included in this study, all older than 18 years. The participants’ answer sheets were identified by unique identification numbers and their identity was not known to the authors, except AN.

Two types of questionnaire were used: first, interview-type questions (S1 File) and, second, a seven-day questionnaire diary (S2 File); both were completed between June and July 2018. The questionnaire and diaries were introduced to the study participants in community meetings led by AN. During the meetings, AN would speak in English and Rukiga as necessary. The completed questionnaire sheets were returned to the CTPH centre and stored in a locked room. In August 2018 data was entered into a Microsoft® Access database by DH and BD, keeping the identity of interviewees anonymous. If clarifications were required these were performed by AN.

### Background questionnaire

The background questionnaire consisted of 29 questions, most of which could be answered by ticking an appropriate box (S1 Data File). Information collected was about age, gender, ethnicity, education, number of people living in the household, marital status and the number of children. Questions including the types and number of latrines used, number of mosquito nets, and any ill health and associated symptoms experienced were also asked. Symptom severity was indicated as mild (*‘does not interfere with your day too much’*), moderate (*‘interferes a lot with your life, but you can be independent’*) or severe (‘*you were unable to behave normally and needed substantial help’*). A request for the vaccination history of each participant and their family members, as per their vaccination records, was included. The final question asked the study participants to identify any animals from a list that they had seen, touched or got close to. The background information was confirmed in a follow-up meeting, which was useful to get the participants ready for completing the diary.

### Diaries

The diary included 49 questions, 48 of which were in a checklist format, allowing participants to answer with a tick (S2 Data File). People were asked to record questions relating to hygiene, such as toileting frequency and location, as well as details around handwashing and food preparation. Behaviours associated with an increased risk of zoonotic pathogen transmission, such as contact with animal faeces or blood, handling raw meat, or being bitten or scratched were included. Questions on daily eating habits included the number of daily meals, food types, treatment of leftovers, and washing of hands and utensils. These questions were designed to capture habits that might lead to increased pathogen exposure as well as nutritional status, that might impact infection susceptibility. Disease symptoms were recorded, including the severity of the symptoms experienced by the participant or of anyone else in the household. Visits to a doctor or traditional healer and details about any medicines taken were recorded.

Participants were also asked if they touched other people, saw other people touching each other, or attended any group meetings that day. The final question asked participants to identify any animal-to-animal contact that they had observed with this question: *“Today have you seen the following touch or got very close (e.g. be together in the same field, trees or plantation)”*. This was done by drawing lines (links) between two identical lists of animals to form an interaction network. Each line then represented that the animals were observed to have close contact (were together in the same field, trees or plantation). Each animal was a node in a pictorial network. The list included 22 taxa: humans, eight domestic (cow, goat, sheep, pig, cat, dog, chicken, rabbit), two peri-domestic considered as one taxa (mouse and rat were summed altogether for the sake of comparison as data from the diary did not include mouse), and twelve wild animal taxa (gorilla, monkey, baboon, bushpig, civet, chimpanzee, elephant, colobus, porcupine, duiker, bushbuck, and squirrel). Questions relating to topics of a sensitive nature such as HIV infection and illegal activities, such as hunting wildlife for food and entering the national park to use its resources, were not asked.

### Ethics approval

Ethics approval was given by the Massey University Human Ethics Committee (Permit number #SOA 17/12, DH), the Royal Dick School of Veterinary Science Human Research Ethical Review Committee (HERC, BD), and Mbarara University of Science and Technology Research Ethics Committee (CTPH, GKZ). No wild, domestic or peri-domestic animals were handled in this study. All participants in the study provided informed consent, and no children under 18 took part in the study.

### Data analyses

#### Demography and health

A descriptive analysis and summary statistics were performed on the data captured from the background questions and the diaries associated with the general health of the community, focusing on reported symptoms and the contacts between species. To check potential correlation between symptoms felt when participants were unwell between background questionnaire week and the diary week we performed a simple Spearman’s correlation test. Then, we present the information on participants who declared contact with gorillas plus timespan of a particular event of direct contact and information relevant for cross-species pathogen transmission.

#### Human social contact data

For direct social contacts, data was collected in age classes and the frequency (0-5 and above contacts per day) for each pairwise interaction. Frequency histograms for contacts between all age class combinations were created. We calculated all human-human direct contacts (counts) and their non-parametric 95% confidence intervals based on a bootstrap procedure with 1000 interactions per unique pair of age classes, containing the interviewee and their direct number of contacts. Then, observed contact rates were calculated as the number of days on which the interaction between age classes was seen (presence or absence) over seven days. While children were not included as participants in our study, we believe asking for observed contacts goes some distance to collect unbiased data about their contact patterns. This means the calculation for observed contact rates (sum of days on which the interaction between age classes were seen for all individuals, separated by age class) is different from the self-declared direct contacts (sum of contacts on for each interaction for all individuals, separated by age class).

#### Human-animal interface

First, data were gathered for each participant, tabulating how many times they had close contact with animals over the week. Then, timeline plots were produced to illustrate the number of self-reported direct contacts (touching) or sightings of animals or their faeces per individual over the week.

Human-animal interface networks were produced from the observations of contacts from two sources: 1) Contacts from the background questionnaire week (link weight = number of participants reporting that contact), 2) Sum of all contacts along the week (link weight = sum of the contacts seen along the week/7). To explore contact information consistency, we correlated the link weights between the background questionnaire week and diary week with Spearman’s correlation test. We believe this approach helps to complement the actual contacts each participant had with contacts that happened in the background questionnaire week and give a broad picture of the local context of contacts. We calculated the animal-animal contact network structure using a network science approach, where a node is a taxon (a species or group of species in several cases, like duiker).and the links between each pair of taxa have weights given by the total number of events along the week when the participants have seen a direct contact or a close contact (be together in the same field, trees or plantation). Where appropriate for network analyses, we converted the edges to binary (present or absent).

To measure how connected the two networks of contacts were we calculated two descriptors: connectance and assortativity. Connectance measures the proportion of binary links observed in a network in relation to the number of potential links it could maximally have. We also quantified the relative frequency of interactions within versus between particular species or groups, using the nominal assortativity metric “r” [67]. We calculated the metric r for two levels: node-node assortativity and assortativity based on broader groups (domestic, peri-domestic, human, wild). This metric is positive when there is a pattern of more contacts with that same node (or group) than with other nodes (groups), and negative when the network presents a disassortative pattern, i.e. when a node (group) has more contacts with other nodes (groups) than with the same. Networks were built in R 4.0.3 [68]; the calculation of metrics and network drawing were performed with the packages *igraph* and *ggplot2* [69,70].

## Results

### Demography and health

#### Background information

All the participants recruited for this study completed the background questions (S1 Data File) and the diaries (S2 Data File), except participant 83, who did not complete the background information. In a small proportion of the questionnaires, individuals did not declare their age (N=2), village (N=1), ethnicity (N=2) or gender (N=1). All participants filled the diaries, but there was variation in the number of days completed: one participant completed the diaries for two days, two people completed them for five days, ten people completed them for six days, 78 participants completed them for seven days, seven participants completed them for eight days, one person completed them for nine days, and one person completed them for ten days. These answers add up to 700 days of daily data for 100 participants.

The sample comprised 81% male to 19% female participants, from 19 to 70 years old (median = 35 years old, S3 Fig). Seventy-five percent of the participants were married, with their age at marriage ranging from 14 years to 30 years with a mean of 21 years (Table 1). Of the 98 respondents that recorded their ethnicity, 93 were from the Bakiga tribe, two from the Banyankole tribe and one each from the Basonga, Batwa and Mukonzo tribes. Half of the respondents resided in Mukono and half in Nkwenda villages. The median number of people living in the same household with each participant was 2, varying from one to 10 people, and the number of children varied with age class of participant (S4 Fig). The background questionnaire included a question that sought to identify diseases that any of the participants had ever suffered from, but unfortunately this question was generally misunderstood, since many identified diseases that they had been previously vaccinated against. Because of that, the answers were not included in the analysis. Most interviewees used pit latrines (98%; 73% had covered pit latrines) and had at least one bed net in the house (Table 1). The mean number of bed nets per household was three. Regarding healthcare behaviour, over half (N=51) of the participants reported feeling unwell in in the week preceding diary completion, reporting at least one symptom (S5 Fig). Thirty participants went to the hospital or health centre to get treatment. Regarding treatment, 9 reported symptoms but did not seek treatment, 10 participants sought self-treatment, and 6 sought traditional treatment with the local healers.

**Table 1.** Background information on Participants in Mukono Parish, Uganda. These questions were answered before the collection of information on diaries, in the previous week. We retrieved information for 99 participants for the background data and 100 participants for the diary data.

| Background information | Value |
| --- | --- |
| Age (mean ±SD) | 37.9 ± 12.6 |
| **Marital Status** |  |
| Married | 77 |
| Not married | 22 |
| Age when married (mean ±SD) | 20.7 ± 3.7 |
| **Education Level** |  |
| Primary complete | 52 |
| Secondary complete | 40 |
| University or College | 20 |
| **Gender** |  |
| Male | 80 |
| Female | 19 |
| **Ethnicity** |  |
| Bakiga | 93 |
| Banyankole | 2 |
| Basonga | 1 |
| Batwa | 1 |
| Mukonzo | 1 |
| Not declared | 1 |
| **Village** |  |
| Mukono | 50 |
| Nkwenda | 49 |
| **Unwell in the last week** |  |
| Yes | 51 |
| No | 48 |
| **Bed Net** |  |
| Yes | 99 |
| **Visited health worker last week** |  |
| No | 69 |
| Yes | 30 |
| **Visited hospital last week** |  |
| No | 79 |
| Yes | 29 |
| **Self-treatment last week** |  |
| No | 65 |
| Yes | 34 |
| **Visited traditional healer last week** |  |
| No | 93 |
| Yes | 6 |

#### Diary information

Regarding meat preparation, 58 of the participants reported in their diaries that they had handled raw meat (N= 101 entries for handling raw meat), 14 had been bitten or scratched by an animal (N=22 entries) and 41 participants had come into contact with the blood of an animal (N=66 entries for touching blood) at least once along the week. All participants reported washing their hands before eating at least once during the week, but did not always use soap to wash their hands, which was done 72% of the time (N=700 entries for all days, N=504 answered “yes” to wash hands with soap before meals). Participants who reported having washed hands with soap before meals and also washed hands with soap after going to the toilet at least once a week summed 94, and using soap for both situations (used soap before meals and after toilet) happened 63% of the time. Regarding nutrition, it was more likely that people would report eating two meals a day (42.3%) than one meal (25.8%) or three meals (25.5%). The staple diet consisted mainly of posho, beans, matooke and porridge with 97.3% of all meals recorded consisting of one or more these carbohydrates. Meals containing animal protein were recorded in 10.7% of all recorded entries.

In the diary, people reported feeling unwell on 281 of the 700 total days recorded (40%) with 11.6% of these records indicating moderate symptoms and 6.4% reflecting severe symptoms. Most participants reported feeling at least one symptom along the week (N=75). In contrast, 48 participants did not report any symptoms during the background survey week. Nevertheless, the number of symptoms felt during the background survey week was positively correlated with the sum of felt symptoms during the diary week (Spearman’s correlation test rho=0.70, p<0.0001). Only 23 participants did not declare any symptom at any point in both data sources (background survey week and diary week).

Of the syndromes reported, headaches and tiredness were the most common (Fig 2). Pain and coughing were reported often (8.7% and 9.5% of the time, respectively), while diarrhoea and fever were less common (2.4% and 4.5% of the time, respectively); itching and skin rashes were reported the least. Two individuals reported seven symptoms on a single day. A person was more likely to feel unwell for one, two or three days (20.2%, 12.9% and 5.1% respectively), compared to four or more days (less than 1%). The average number of symptoms reported was 1.7 per person (min=0, max=7 symptoms per day), where it was most common to report no symptom (59.8%), or 1 symptom per day (20% of entries).

**Figure 2.** For a week, symptoms and their severity were reported in participants diaries in Buhoma, southwestern Uganda (2018).

People were more likely to pass a bowel movement more than twice in a day compared to once a day (73.3% versus 26.7%), which may reflect a high incidence of gastrointestinal disease in the human population. Despite the frequent number of bowel movements reported per day (mean=2.05, median=2, max=5), people only acknowledged having diarrhoea in 17 entries of the diary reported by nine participants. The diary information showed that forty-six people sought medical help from a clinic and took medication with a further 14 people seeking advice and treatments from a traditional healer along the week, varying from one to seven visits along the week. From the people who visited traditional healer, 10 ended up taking medicine from the traditional healer; 23 participants reported taking medicine they prepared at home along the week.

### Human social contact data

We report self-reported close contacts (touching) for 100 individuals (Table 1, Fig 3). Participants reported a total of 5777 direct contacts, but 113 contacts from 2 participants were removed because we could not retrieve the participants’ ages. Thus, we ended up with 5664 human-human direct interactions. Moreover, participants reported 2491 observations of interactions between other people from various ages (when they saw people touching other people), adding up to a total of 8155 human-human interactions in our study. Age classes with highest observed contacts were interviewees of age class 41 or more who interacted with people aged 21-40 (S6 Fig). Self-reported direct social contacts varied from an average of 0.76 to 2.30 contacts per person per day for any age class (Fig 3, S1 Table). Observed contacts varied from an average of 0.65 to 3.53 (mean= 1.66 average number of days of contact per age class, S6 Fig).

**Figure 3.** **Frequency histograms with self-reported direct human-human (N= 5664) contacts around BNP.** Data represents age classes for 98 individuals (there was no age information for two individuals) and the number of people they touched along one week in a self-reported diary. No children participated in the questionnaire.

### Human-animal interface

From the diary questions, where participants were asked if they had seen a certain type of animal or their faeces that week, the domestic animals such as chickens, goats, cows and pigs were reported as the most common animals seen compared to wild animals. Peri-domestic animals were also frequently seen. Seeing was more common than touching, and wild animals were rarely reported to be touched (Fig 4). The animals that were reported to be touched most frequently were goats, chicken, pigs and cows, cats and dogs. However, there were very rare records of touching wild animals. One person reported to touch a civet and a monkey at the same day. One participant reported to have touched a duiker, and there were two events of touching a squirrel. Moreover, 4 participants reported having touched a monkey, with one participant reporting having touched a monkey twice in a week (S3 Table). Contact with animal faeces/dung—only reported for domestic and peri-domestic animals, since their faeces are more easily recognizable—are reported in S7 Fig. Contact with the faeces of different animals followed the trend for direct contacts reported, with sightings being more common than direct contacts—touching the dung of chicken, goats, cow and pigs were the most common direct contacts reported.

**Figure 4.** **Sightings and direct contacts between participants, wildlife, domestic and peri domestic animals reported by ordered participants along a week by self-reported dairies in Buhoma, Uganda.** A. Sightings in dark gray B. Touching directly the animal in dark gray.

Over the seven-day period that the diaries were recorded 20 participants reported seeing mountain gorilla at 40 occasions. On 21 of those occasions, the participants ate two or less meals the day of sighting, in 29 cases they had more than one bowel movement in a day, with 2 individuals reaching 4 bowel movements in a day, and in 10 of the 40 occasions of gorilla sightings, the participants drank unboiled water from sources. From the 20 individuals who declared seeing a gorilla at 40 occasions, a majority of people felt unwell (31/40 occasions), and there were 23 occasions when they reported that others in the same household were unwell on the same day the participant saw a gorilla. The average number of symptoms of people who saw a gorilla reported was 1.52 (median=1, min=0, max=6), slightly higher than the average number of symptoms per person along the week was 1.36 (median=1, min=0, max=7). The average number of symptoms per person for those occasions when the participants did not see a gorilla was 1.25 (median=1, min=0, max=7). The most common symptom of those who saw a gorilla was tiredness and headaches but coughing and fever were reported too. In one occasion, a participant reported a severe cough on the same day of a gorilla sighting. Respondents recorded between one and five gorilla sightings over the week. Nine people reported multiple gorilla sightings over the week. On one occasion there was physical contact between gorilla and a human (S8 Figure). At the same day of this physical contact, the participant touched other animals (cow, chicken, goat) and saw several other animals, including monkeys and baboons. This participant also had four bowel movements that day and declared having respiratory symptoms (coughing) two days before touching the gorilla and mild pain at the day of contact. This individual reported coughing and headache symptoms in members of his household three days after the day of contact.

The contact networks illustrate that people involved in this study had regular contact or had been in close proximity to both domestic animals and wildlife (Fig 5). Both networks were moderately connected; The connectance value for the human-animal contact network using the diary data (divided by 7) was 0.31, and 0.26 for the background questionnaire week network. The observations of contacts revealed many close contacts between groups of humans, domestic, peri-domestic and wildlife. Regarding assortativity (preferential links) in the network, taxa tended to contact their similar taxa over other taxa (r=0.24 for diary network, r=0.29 for the background questionnaire week network). In addition, the same group (groups = domestic, human, peri-domestic or wildlife) also tended to contact the same group over contacting other groups (r=0.40 for diary network, r=0.32 for background questionnaire week network). The complete interaction list for the contact networks considering background questionnaire week interactions and summed diary interactions is available in S3 Table. From the close contact networks, the most common interaction observed between groups were cow-person (S3 Table), followed by goat-person, chicken (domestic)-person, followed by pig-person. Person-person contacts made up for the highest number of interactions, followed by goat-goat, cow-cow and chicken-chicken. Peri-domestic contacts were the 12th most common interaction, while the wildlife species with most reported interaction events were primates. The most reported interaction between wildlife taxa in close contact with humans were baboons, then gorillas, then monkeys. Most common wildlife-wildlife in-taxon contacts were gorilla-gorilla (N=56) and monkey-monkey (N=56). Wildlife-domestic close contacts were rarely seen, such as squirrel-rabbit, gorilla-cow, chimpanzee-dog, or elephant-dog. Person-gorilla contacts were the 36^th^ most common observed interaction in the network. The complete list containing the number of taxa each animal interacted with (including themselves) is available in S4 Table.

**Figure 5.** **Local close contact networks reported in Buhoma, Uganda (2018).** Data was collected from self-reported surveys. Node size and link line width are proportional to number of interactions at taxa-level (number of taxa the node interacted with) and link-level (value of self-reported and observed contacts summed), respectively. Colours illustrate people in yellow, wildlife in red, peri-domestics in green, and domestic animals in purple. Animal-to-animal data comes from observations of people who reported seeing an animal touching or getting very close to another animal. A. Seen last week (background questionnaire week), B. Seen during the week of Diary completion. C. Correlation between matching pairwise interactions. Labels in gray represent interaction of nodes from different groups.

During the diary completion week, humans interacted with 18 taxa, followed by dogs and cows, which were seen in close contact with 13 and 10 taxa. The wild taxon in close contact with the smallest number of other taxa was the civet (1 taxa), followed by duiker and porcupines (all interacting 2 taxa or less, including themselves). Gorillas had rarely been observed in close proximity or were seen touching domestic animals (dogs and cows) and wildlife (baboon and monkey).

## Discussion

Our study reveals that the human population of the Buhoma sector around BINP is generally suffering from poor health and lives at the interface of substantial domestic, peri-domestic, and wild animal interactions, with important implications for the risk of cross-species pathogen transmission events. Examining contact patterns and community health is important to understand those risks and contribute to initiatives that increase human and ecological health. Over half of the participants reported symptoms of disease within the previous week with varying levels of severity. Contacts with domestic and peri-domestic animals were very common, and human-human contacts were the most common contacts reported, as expected. Seeing wild animals was much more common than touching them, with only nine directly reported events of touching a wild animal, and one case of person touching a gorilla. These numbers make up much less than 1% of total animal-person direct self-reported contacts (N=1424), with most direct human-animal contacts being with domestic or peri-domestic animals. However, people’s perceptions of close contacts along the week revealed hundreds of close contacts between humans and 20 animal species considered in the survey (S3 Table, S4 Table). Below we discuss the self-reported health patterns, focusing on the interaction between people and mountain gorillas, the local context, social contact patterns and the broad human-animal interface. Then, we discuss caveats of this study and future directions.

### Demography and health

People declared feeling unwell, feeling at least one symptom 40% of the time along the week, an indicator of poor health, with headaches and tiredness being the most reported symptoms. These could be related to the general lack of good nutrition, inadequate intake of water or other fluids, or may reflect underlying disease—both of which could result in a compromised immune system. The frequent number of daily bowel motions of study participants could indicate underlying gastrointestinal disease. With a local human prevalence of intestinal parasitism ranging from 25% to 57% and the prevalence of other enteric pathogens at 49%, including *Giardia* and *Cryptosporidium* [71], it is possible that diarrhoea was under-reported in this study (17 cases in 700 entries, 2.4%), given that people reported passing a bowel motion up to five times in a day without reporting diarrhoea. In addition, over a quarter of the participants reported drinking un-boiled water, potentially exposing them to water-borne disease risk.

Regarding disease symptoms, although guidance was given in the questionnaire regarding severity levels, all information gathered was self-reported and therefore subjective. Moreover, if the ‘normal’ state for an individual is feeling constantly tired with headaches (Fig 2), possibly due to poor nutrition, heat and dehydration, the individual may not report these as symptoms. This may also be the case for individuals suffering from chronic diseases who have lived with the symptoms for many years and have learnt to tolerate them. This may mean that the symptoms’ severity could be under-reported so these results should be interpreted with caution, such as the difference between reported diarrhoea and the frequency of defecation reported. It is common for people in this community to only eat two meals a day, with a diet that is high in carbohydrates, low in protein, and generally of low nutritive value, suggesting that most of the local population is likely to be malnourished. Food insecurity in the area was beyond the focus of this study and should be assessed in depth using survey methods designed for that purpose (e.g., [72]).

Considering risk factors for disease exposure, 68% of people reported handling raw meat and animal blood, and 28 people indicated that they had cuts on their hands during the week. Eleven participants had contact with animal blood and had cuts on their hands on the same day. These events are associated with an increased risk of the transmission of zoonotic diseases. Several studies in the area surrounding BINP and other parks have demonstrated the sharing of pathogens between the local human population, domestic livestock and wild animals, many of which could be transmitted by these high-risk activities [73–77]. The use of pit latrines also increases the risk of disease [78,79]. Together, these findings may explain the high level of disease symptoms reported in this study. All participants provided stool samples that may reveal underlying disease and will be analysed in future studies. A community with limited access to health care of good quality and with a high density of malnourished and immunocompromised people can act as a potential reservoir for pathogens that could be transmitted to non-human great apes such as the endangered mountain gorilla. Although many people reported seeking medical help, the quality of care they received is unknown. The relatively high level of handwashing reported by the participants is likely due to the regular workshops run by CTPH throughout the local communities using a network of Village Health and Conservation Teams (VHCTs), community health workers also trained to promote biodiversity conservation. A focus on preventative human health care, like the CTPH workshops, immunisation programs and surveillance should be prioritised for the local communities bordering the BINP. Commonly reported diseases for this region and other rural parts of Uganda include malaria, measles, intestinal parasites, typhoid and skin disease [80]. For those, fever, coughing and respiratory distress the most reported symptoms, whereas the symptoms reported in our survey could be associated with malaria and the other diseases mentioned. Moreover, several zoonoses including leptospirosis, Q fever and rickettsial fevers occur locally [6,81]. For privacy reasons, people were not asked about their HIV status. However, it could be assumed to be around the values of prevalence for southwestern Uganda (7.9%, [82]).

### Human social contact data

Data on human contact patterns are scarce in Uganda [51], and gathering sufficient data about social contacts can help understand the spread of infectious diseases. Our results show that reported contacts among humans were numerous—the participants declared to be interacting with five people or more (regardless of age class) in 71% of the days, with a mean of at least 8 contacts per day and a median of seven contacts per day. This pattern of numerous interactions per individual per day was reported in another study recently conducted in southwestern Uganda [51], which reported an average routine contact of 7.2 individuals per participant. It is interesting to see this convergence of results between self-reported assessments guided through workshops in our study and a study based on interviews under a similar social context and region [51]. Self-reported contacts were restricted to individuals of 18 years old or more, and thus are limited by the age groupings. With the contact data is biased towards adults, this allows only limited inferences about child contacts and older adults due to how the data for age classes were divided, aggregating all ages from 41 years or older. Observations of contacts, reported in the diaries, also pointed to more contact events for the ages of 21 and older. However, these data give more insights into contacts for children and teenagers because, while participants were always older than 18 years old, they could observe the presence of a variety of age classes interacting during their daily routine (S6 Figure).

### Human-animal interface

Systematic data on contact between humans and animals and between different animal taxa are relatively rare but can be valuable for understanding the complex process of how pathogens spread from contact with other animals or their environment [38]. As expected, most reported and observed contacts occurred between people and domestic animals, emphasizing the importance of livestock in the area and how they are deeply associated with social and economic relationships [40]. Contact (and thus pathogen transmission) is likely to be more frequent between domestic mammals and their wild relatives than between humans and wild primates [83]. The potential of pathogen transmission through taxa and their environment should be a priority in future studies in this area. For instance, in this area, a recent study detected *Giardia* in 5.5% of livestock in this area that has an active human-livestock-wildlife interface [84].

The networks of close contacts plus direct reports of contacts revealed hundreds of interaction events, with a positive correlation (Spearman’s correlation test rho=0.67, p<0.0001) between the interactions reported for the background questionnaire week and the interactions occurring during the diary week (Fig 5). From those, despite rare events, wild animals were mentioned to be having close contacts with humans, for instance, gorillas, baboons, squirrel, monkey, and duiker (S3 Table, S4 Table). One of the participants declared direct contact with a civet, and one participant declared having direct contact with a gorilla. The participant who touched a gorilla was male, Bakiga, and had cuts on his hands before and after the day of contact, in addition to symptoms of concern. He had contact with people on the same day of the contact with a gorilla (day 4). He declared to have touched chicken, cow, and goats on the same day, and also touched the dung of chicken, cow, goat, pig, and rat at the same day he touched a gorilla. At that time, he was part of the human-gorilla conflict resolution teams (HUGOs) and declared having headache and respiratory symptoms (coughing) on day 7 in his household members three days after touching a gorilla. The participant declared a visit to the clinic on day 2 of the diary. Nevertheless, of concern are the respiratory symptoms of coughing and breathing difficulty, given that respiratory viruses can cause severe disease and lead to fatalities in gorilla populations [7,18,85]. This participant reported coughing symptoms two days before the contact. Given how contagious COVID-19 is and having spread from an asymptomatic keeper to eight captive gorillas in San Diego Zoo Safari Park [86], this study provides useful information to minimise human and gorilla disease transmission during the pandemic.

Gorilla sightings appear to be frequent as over half the respondents had ever seen a gorilla, and nine of these reporting multiple gorilla sightings over the week, which is similar to the results of a previous study in this area [71]. Several of these respondents also reported seeing other wildlife such as monkeys and baboons, which may indicate a group of people that may work near the park. A group likely to report a high number of gorilla sightings are the human and gorilla conflict resolution volunteers (gorilla guardians or HUGOs). HUGOs should be prioritized for preventive health measures in the region; it would be interesting to measure contacts more intensively for this group, composed of people responsible for moving mountain gorillas who have ventured onto agricultural land backing onto the forest. Moreover, there were 1.57 reported gorilla-person contacts/day reported (and 3 reports for background questionnaire week) and 5.57 gorilla-gorilla contacts/day (10 records for the background questionnaire week network), which brings an interesting perspective on observer bias and on how frequent this contact is. When a person gets close to a gorilla, it is highly likely that this person also sees a gorilla in close contact with other gorillas.

This study has revealed species within the Bwindi Impenetrable Forest ecosystem that are in close contact. As other studies have indicated, humans have cross-species contacts [58] with gorillas, which are to be expected given the local mountain gorilla conservation programs and tourism. Also, some gorilla families have home ranges close to the park’s perimeter, and that move out of the forest, spending a considerable amount of time close to cultivated agricultural areas [64]. This may explain the rare events of sightings of cows and gorillas (0.14 contacts/day, zero for the background questionnaire week) and dogs and gorillas (also 0.14 contacts/day and zero for the background questionnaire week, S3 Table) reported. We need to gain a better understanding of where this interface may be most frequently active. This will help determine the strategies that could be used to minimise the risk of disease transmission at this interface, in the absence of wider change in ultimate drivers. Goats have free range close to the border of the forest, and this hard edge is probably where goats and gorillas are more likely to spatially overlap and to share waterways and grazing areas. Cows are more likely to be tethered at night but like the goats are walked to new grazing areas each day.

Although humans and great apes are more distantly related to domestic animals, this does not preclude the transmission of pathogens, particularly those that can infect multiple hosts, from goats and cattle to humans or wild great apes [83]. Monkeys and baboons were reported as seen near the gorillas, a finding reported previously [87]. Baboons and gorillas share several intestinal parasites including *Cryptosporidium, Giardia, Trichuris* and *Strongyloides* species [87]. It is likely that gorillas and monkeys have pathogens in common due to their spatial overlap and shared habitat. Respiratory viruses with possible transmission between humans and baboons [88] and humans and New World monkeys [89] have been observed. In our network, baboons, monkeys and gorillas had similar interactions among themselves (Figure 5, S3 Table). As well as being potential victims of disease, this highlights the possible risk of monkeys or baboons acting as reservoirs or intermediate hosts for diseases that affect both humans and gorillas.

Despite intense action of different actors in the region to promote conservation and public health, including a One Health national agenda [90], empirical studies need to quantify these relationships between human beings and wildlife more thoroughly. Though we specifically did not ask the nature of any contacts, our study is limited to what participants were comfortable reporting. Activities related to the collection of resources in the park are regulated via permits for multiple use zones in parishes bordering the park, and outside of these areas, law enforcement can be applied. Pressures on living conditions and cultural practices still lead to unauthorized access to the park’s resources, such as collecting herbs, honey, and wood. We acknowledge that hunting and illegal poaching still happens in the region, including recent cases of gorilla killings [91], which is negative from a biological conservation perspective, but also a complex issue that needs to be further addressed since there is a high subsistence demand in the region [92,93].

### Limitations of Study

This study has significant limitations, such as recall bias, typically associated with diary-based research [94]. However, filling diaries for a short period tends to minimize recall bias due to the small-time lag between an event occurring and being recorded compared to a questionnaire [95]. Another bias is the non-random nature of the recruitment of participants since their selection was based on literacy and age. Although the questionnaire and diary questions asked about the health of all household members, the ages of any unwell members were not captured. Because of these limitations, it is possible that the frequency of disease symptoms reported in the diaries in this study underrepresents the actual disease in the community. Moreover, literacy can be associated with the tasks that a person might do in their day-to-day lives, with low literacy levels associated with more prolonged episodes of sickness [96]. More literate adults are more likely to work outside of the home in occupations such as teaching, nursing, or running their own business, whereas nonliterate individuals are more likely to be doing manual work at home and working with animals and crops [3]. The latter activities present a higher risk for human contact with livestock (which is the most common type of cross-species contact, see S3 Table) or wildlife and associated pathogen transmission. We will investigate community activities in future works but note that most activities in the area point to subsistence farming [92,97], which is noticeable in our results showing the frequent contacts with livestock.

Furthermore, male participants were overrepresented in this study (81% males, 19% females). This disparity in the selection of study recruits may partly reflect convenience, but partly because of gender biases and patriarchal systems, meaning more men volunteer as our community member who administered the questionnaires (AN) was male. As different genders can perform different activities, the diseases associated with these tasks can be gender-sensitive [98]. An example of this is the gender-specific division of labour, which is a feature of the Bakiga tribe, with men responsible for hunting and building houses and women collecting firewood and fetching water [99]. Future analysis should be more representative of the population in terms of participants’ gender, literacy level and ethnicity. This is especially important for marginalised groups, such as the Batwa (only 1 participant in our study), who were traditionally hunter-gatherers and forest dwellers and who have a low literacy level [100]. The Batwa were resettled from within BINP when the park was established, and they have been defined as refugees in the villages at the forest margins [24], with privatised land surrounding the forest largely owned by the Bakiga people (the majority of participants, N=93). Despite local actions of NGOs and institutional initiatives, the edge-dwellers’ situation remains critical in terms of living conditions and well-being [101], and the survival of the Batwa in the future is threatened. Furthermore, many Batwa living in extreme poverty are forced to use public land resources, thus increasing their exposure to diseases from polluted groundwater and latrines [24]. Strategies, such as poverty alleviation, addressing crop raiding and illegal access to the park, improving revenue sharing, and local governance are envisioned for this area at governmental levels [102]. This region demands long-term efforts to combat poverty, discrimination against minorities, and tailoring of interventions to their needs.

### Future studies for managing disease risk

Acknowledging the complexity of the interactions between humans and animals and human behaviour are essential steps for understanding systems, including developing infectious disease dynamic models and for meaningful interventions to reduce disease risk [38,103]. Human and animal dimensions are entangled in the Buhoma sector, where people’s livelihoods are connected with BINP due to the proximity to the park. Concurrent research from our group investigates infection prevalence and cross-species transmission events, using next-generation sequencing metagenomic studies from faeces collected from cattle, goats, humans, and gorillas in the BINP region. The results from these studies will assist with informing future work measuring relevant components for cross-species transmission. To manage the risk of disease transmission in the region, further detail is needed regarding non-human great apes that are in close contact with humans and domestic species. Future studies may look at the spatial and temporal overlap of all these species, as well as specific activities associated with a high risk of pathogen transmission. Detailed mapping of people and animal movements can reveal spatial or temporal overlaps between species. A thorough medical survey, including biological sampling of individuals who are in regular contact with the gorilla groups, may highlight subgroups that present a higher risk of disease transmission. These subgroups include visiting tourists, researchers, gorilla trackers, children, park rangers, ethnic groups, and the HUGOs. In addition, future studies could do more in-depth interviews and participant observation with the people that reported regular sightings of wildlife to gain more insights into the high-risk activities and social determinants that could lead to the transmission of pathogens and to an understanding of the wider political economy and livelihood challenges regionally. A preventative health program to manage intestinal parasites and manage livestock to reduce contamination of shared waterways should be considered in goat flocks and cattle herds close to BINP. Preventative health programs and disease surveillance of the local human and gorilla populations should continue to prioritize improving livelihoods and conservation in the area. Moreover, disease surveillance of tourists entering the park could further inform the disease risks within the local ecosystem. The One Health Initiative organized by CTPH, with support from the Uganda Ministry of Health and other organizations, led to an increase in family-planning users, support for conservation, and reduction in incidence of gorilla disease, including giardiasis and scabies since the creation of CTPH in 2003 [34,84].

## Conclusion

This study highlights a local human population that is generally unwell and may act as a reservoir for pathogens. Humans and domestic animals form a tightly coupled system that may act as a reservoir for infectious diseases that could spill over to endangered wildlife, including the mountain gorilla. Despite having several caveats, our findings are relevant to inform local community workers and stakeholders of the overall health condition and contact patterns of the local community around BINP. Several behaviours that may increase zoonotic disease transmission risk were identified for Mukono and Nkwenda villages in the Buhoma sector. With fewer than 1000 mountain gorilla mature individuals remaining in their natural habitat [15], it is a matter of urgency that we gain the knowledge that will allow us to best mitigate the risks and prevent the transmission of infectious diseases to this endangered species. More broadly, the improvement of healthcare and living conditions in the region is of utmost importance to reduce the burden of pathogens in humans and wildlife. In this case study, we provide evidence of an active interface between humans and other species, including cattle, goats, baboons, and the endangered mountain gorilla around the BINP. We offer the first characterization of the human-animal contact network around BINP based on survey data and report frequent interactions indicating strong potential for pathogen cross-species transmission among various animal species.

## Acknowledgements

We thank the team from CTPH in Uganda for their invaluable services to the community. We were assisted by Richard Bagenyi and Stephen Rubanga in the local administration, planning and executing of the project. We thank Janelle Wierenga for the fruitful discussions about BINP and Neil Anderson for the review of BD’s MSc project.

## Supporting Information

**S1 File. “File Background questions.pdf”** Virtual version of the questionnaire that was printed to the participants in Buhoma, Uganda (2018).

**S2 File. “S2 File Diary questions.pdf”.** Virtual version of the dairy that was printed to the participants in Buhoma, Uganda (2018).

**S1 Data file. “Data_File_S1_Survey.xlsx”.** Relevant data from the questionnaire containing the answers from to the participants in Buhoma, Uganda (2018).

**S2 Data file. “Data_File_S2_Daily.xlsx”.** Relevant data from the dairy containing the answers from to the participants in Buhoma, Uganda (2018).

**S1 Figure: Photos from Buhoma, Uganda, showing elements that are part of the local living conditions around Bwindi Impenetrable National Park.** A. Set for washing hands and water usage outside a house. B. Water supply in plastic containers. C. Water supply from artesian well. D. Local butchery entrance. D. and E. goats very close to the Park border. F. Cooking area in one of the village’s houses. Photos: D. Hayman.

**S2 Figure. Pictures from Buhoma, Uganda, showing parts of the village.** A. Panoramic view of the street. B. View of the village close to the Bwindi Impenetrable National Park and livestock in the background. C. Closer view of one of the local houses. Photos: D. Hayman.

**S3 Figure. Gender and age distribution of the participants.** Gender bias in our sampling is a significant study limitation.

**S4 Figure. Average number of children per participant age group.** The average number of children per age group of participants varied from no children older than six years old for younger participants (between 16-20 years old) to more than three children for participants above 30 years old.

**S5 Fig. Health care behaviours in Buhoma during a self-reported survey conducted in 2018.** Treatment choices of those who declared to be unwell or seek treatment (abbreviated as “treat.”) during the week before diary completion.

**S6 Figure. Self-reported observed human-human contacts around BNP.** Observed contacts were collected along one week in a self-reported diary per age class.

**S7 Figure. Self-reported direct contacts between humans and animal faeces around BNP.** Contacts were collected along one week in a self-reported diary. A. Sightings of dung in gray. B. Direct contact with dung in gray.

**S8 Figure. Summary of self-reported information for the participant who declared to have touched a gorilla.** Timeline with symptoms reported for a participant who reported touching a mountain gorilla. Sources for the gorilla and hand sillhouettes: Creazilla (CC BY 4.0): https://creativecommons.org/licenses/by/4.0/deed.pt; https://creazilla.com/nodes/832270-two-hands-reaching-silhouette

**S1 Table. Age classes of self-reported direct contacts (n=98) with the people they touched along seven days in Buhoma, Uganda.** Two individuals from the total sample (n=100) did not declare their age and were removed from the analysis.

**S2 Table. Self-reported events of actual touching certain animals or their dung the previous week in Buhoma, Uganda (2018), according to 100 participants.** Values for all species in the questionnaire are shown from highest to lowest.

**S3 Table. Link list containing the events of perception of local close contacts during a one-week survey in Buhoma, Uganda (2018).** The number of contact events is in decrescent order based on the diary week sum of events divided by seven days.

**S4 Table. Taxon-level metrics reported in Buhoma according to the participant**’**s observations of human and animal close contacts**. Data is arranged in ascending order of the number of species a node interacted with, including the node itself.

## References

1. Andersen KG, Rambaut A, Lipkin WI, Holmes EC, Garry RF. The proximal origin of SARS-CoV-2. Nature Medicine. 2020;26: 450–452. doi:10.1038/s41591-020-0820-9

2. Campos Tisovec-Dufner K, Teixeira L, Marin G de L, Coudel E, Morsello C, Pardini R. Intention of preserving forest remnants among landowners in the Atlantic Forest: The role of the ecological context via ecosystem services. People and Nature. 2019;1: 533–547. doi:10.1002/pan3.10051

3. Leach M, Bett B, Said M, Bukachi S, Sang R, Anderson N, et al. Local disease – ecosystem – livelihood dynamics: Reflections from comparative case studies in Africa. Philosophical Transactions of the Royal Society B: Biological Sciences. 2017;372. doi:10.1098/rstb.2016.0163

4. Cilliers S, Cilliers J, Lubbe R, Siebert S. Ecosystem services of urban green spaces in African countries — perspectives and challenges. Urban Ecosyst. 2013;16: 681–702. doi:10.1007/s11252-012-0254-3

5. Mpimbaza A, Walemwa R, Kapisi J, Sserwanga A, Namuganga JF, Kisambira Y, et al. The age-specific incidence of hospitalized paediatric malaria in Uganda. BMC Infectious Diseases. 2020;20: 1–12. doi:10.1186/s12879-020-05215-z

6. Dreyfus A, Dyal JW, Pearson R, Kankya C, Kajura C, Alinaitwe L, et al. Leptospira Seroprevalence and Risk Factors in Health Centre Patients in Hoima District, Western Uganda. PLoS Neglected Tropical Diseases. 2016;e0004858: 1–14. doi:10.5061/dryad.6ns6p.Funding

7. Devaux CA, Mediannikov O, Medkour H, Raoult D. Infectious Disease Risk Across the Growing Human-Non Human Primate Interface: A Review of the Evidence. Frontiers in Public Health. 2019;7: 1–22. doi:10.3389/fpubh.2019.00305

8. de Vos JM, Joppa LN, Gittleman JL, Stephens PR, Pimm SL. Estimating the normal background rate of species extinction. Conservation Biology. 2015;29: 452–462. doi:10.1111/cobi.12380

9. Bellard C, Cassey P, Blackburn TM. Alien species as a driver of recent extinctions. Biology Letters. 2016;12. doi:10.1098/rsbl.2015.0623

10. Thomas JA, Telfer MG, Roy DB, Preston CD, Greenwood JJD, Asher J, et al. Comparative Losses of British Butterflies, Birds, and Plants and the Global Extinction Crisis. Science. 2004;303: 1879–1881. doi:10.1126/science.1095046

11. Rhyan JC, Spraker TR. Emergence of diseases from wildlife reservoirs. Veterinary Pathology. 2010;47: 34–39. doi:10.1177/0300985809354466

12. Aguirre AA, Tabor GM. Global factors driving emerging infectious diseases: Impact on wildlife populations. Annals of the New York Academy of Sciences. 2008;1149: 1–3. doi:10.1196/annals.1428.052

13. Roelke-Parker ME, Munson L, Packer C, Kock R, Cleaveland S, Carpenter M, et al. A canine distemper virus epidemic in Serengeti lions (Panthera leo). Nature. 1996;379: 441–445. doi:10.1038/379441a0

14. Marino J, Sillero-Zubiri C, Deressa A, Bedin E, Bitewa A, Lema F, et al. Rabies and distemper outbreaks in smallest ethiopian wolf population. Emerging Infectious Diseases. 2017;23: 2102–2104. doi:10.3201/eid2312.170893

15. Hickey JR, Basabose A, Gilardi KV, Greer D, Nampindo S, Robbins MM, et al. Gorilla beringei ssp. beringei. The IUCN Red List of Threatened Species 2018. The IUCN Red List of Threatened Species 2018. 2018;8235: e.T39999A17989719. Available: https://dx.doi.org/10.2305/IUCN.UK.2018-2.RLTS.T39999A17989719.en

16. Davies TJ, Pedersen AB. Phylogeny and geography predict pathogen community similarity in wild primates and humans. Proceedings of the Royal Society B: Biological Sciences. 2008;275: 1695–1701. doi:10.1098/rspb.2008.0284

17. Travis DA, Lonsorf E v., Gillespie TR. The grand challenge of great ape health and conservation in the anthropocene. American Journal of Primatology. 2017;80: 1–5. doi:10.1002/ajp.22717

18. Mazet JAK, Genovese BN, Harris LA, Cranfield M, Noheri JB, Kinani JF, et al. Human Respiratory Syncytial Virus Detected in Mountain Gorilla Respiratory Outbreaks. EcoHealth. 2020. doi:10.1007/s10393-020-01506-8

19. Bas Huijbregts, Pauwel De Wachter, Louis Sosthe’ ne NO and MEA. Ebola and the decline of gorilla (Gorilla gorilla) and chimpanzee (Pan troglodytes) populations in Minkebe Forest, north-eastern Gabon. Oryx. 2002;37: 1–6. doi:10.1017/S0030605302000000

20. Rulli MC, Santini M, Hayman DTS, D’Odorico P. The nexus between forest fragmentation in Africa and Ebola virus disease outbreaks. Scientific Reports. 2017;7: 1–8. doi:10.1038/srep41613

21. Wilkinson DA, Marshall JC, French NP, Hayman DTS. Habitat fragmentation, biodiversity loss and the risk of novel infectious disease emergence. Journal of the Royal Society Interface. 2018;15. doi:10.1098/rsif.2018.0403

22. Economics T. Uganda - Rural Population Density (rural Population Per Sq. Km Of Arable Land). 2011 [cited 25 May 2020]. Available: https://tradingeconomics.com/uganda/rural-population-density-rural-population-per-sq-km-of-arable-land-wb-data.html%0D

23. United Nations, Department of Economic and Social Affairs PDivision. World Population Prospects: The 2019 Revision. 2019 [cited 2 Dec 2020] p. 1. Available: https://population.un.org/wpp/

24. Banbury C, Herkenhoff L, Subrahmanyan S. Understanding Different Types of Subsistence Economies: The Case of the Batwa of Buhoma, Uganda. Journal of Macromarketing. 2015;35: 243–256. doi:10.1177/0276146714528954

25. Dunay E, Apakupakul K, Leard S, Palmer JL, Deem SL. Pathogen Transmission from Humans to Great Apes is a Growing Threat to Primate Conservation. EcoHealth. 2018;15: 148–162. doi:10.1007/s10393-017-1306-1

26. Xue Y, Prado-Martinez J, Sudmant PH, Narasimhan V, Ayub Q, Szpak M, et al. Mountain gorilla genomes reveal the impact of long-term population decline and inbreeding. Science. 2015;348: 242–245. doi:10.1126/science.aaa3952

27. Woodford MH, Butynski TM, Karesh WB. Habituating the great apes: The disease risks. Oryx. 2002;36: 153–160. doi:10.1017/S0030605302000224

28. Palacios G, Lowenstine LJ, Cranfi MR, Gilardi KVK, Lukasik-Braum M, Kinani J-F, et al. Human Metapneumovirus Infection in Wild Mountain Gorillas, Rwanda. Emerging Infectious Diseases. 2011;17: 711–713. doi:10.3201/eid1704100883

29. Grützmacher KS, Köndgen S, Keil V, Todd A, Feistner A, Herbinger I, et al. Codetection of Respiratory Syncytial Virus in Habituated Wild Western Lowland Gorillas and Humans During a Respiratory Disease Outbreak. EcoHealth. 2016;13: 499–510. doi:10.1007/s10393-016-1144-6

30. Gilardi KVK, Oxford KL, Gardner-Roberts D, Kinani JF, Spelman L, Barry PA, et al. Human Herpes Simplex Virus Type 1 In Confiscated Gorilla. Emerging Infectious Diseases. 2014;20: 1883–1886. doi:10.3201/eid2011.140075

31. Graczyk TK, Mudakikwa AB, Cranfield MR, Eilenberger U. Hyperkeratotic mange caused by Sarcoptes scabiei (Acariformes: Sarcoptidae) in juvenile human-habituated mountain gorillas (Gorilla gorilla beringei). Parasitology Research. 2001;87: 1024–1028. doi:10.1007/s004360100489

32. Salzer JS, Rwego IB, Goldberg TL, Kuhlenschmidt MS, Gillespie TR. Giardia sp. and Cryptosporidium sp. Infections in Primates in Fragmented and Undisturbed Forest in Western Uganda. Journal of Parasitology. 2007;93: 439–440. doi:10.1645/ge-970r1.1

33. Wolf TM, Sreevatsan S, Travis D, Mugisha L, Singer RS. The risk of tuberculosis transmission to free-ranging great apes. American Journal of Primatology. 2014;76: 2–13. doi:10.1002/ajp.22197

34. Kalema-Zikusoka G, Byonanebye J. Scaling up a one-health model of conservation through public health: experiences in Uganda and the Democratic Republic of the Congo. The Lancet Global Health. 2019;7: S34. doi:10.1016/s2214-109x(19)30119-6

35. Plantier JC, Leoz M, Dickerson JE, de Oliveira F, Cordonnier F, Lemée V, et al. A new human immunodeficiency virus derived from gorillas. Nature Medicine. 2009;15: 871–872. doi:10.1038/nm.2016

36. Keele BF, Heuverswyn F van, Li Y, Bailes E, Takehisa J, Santiago ML, et al. NIH Public Access. 2008;313: 523–526.

37. Lloyd-Smith JO, George D, Pepin KM, Pitzer VE, Pulliam JRC, Dobson AP, et al. Epidemic Dynamics at the Interface, Human-animal. Science. 2009;326: 1362–1368. doi:10.1126/science.1177345.Epidemic

38. Plowright RK, Parrish CR, McCallum H, Hudson PJ, Ko AI, Graham AL, et al. Pathways to zoonotic spillover. Nature Reviews Microbiology. 2017;15: 502–510. doi:10.1038/nrmicro.2017.45

39. Wolf M. Is there really such a thing as “one health”? Thinking about a more than human world from the perspective of cultural anthropology. Social Science and Medicine. 2015;129: 5–11. doi:10.1016/j.socscimed.2014.06.018

40. MacGregor H, Waldman L. Views from many worlds: Unsettling categories in interdisciplinary research on endemic zoonotic diseases. Philosophical Transactions of the Royal Society B: Biological Sciences. 2017;372: 1–9. doi:10.1098/rstb.2016.0170

41. lo Iacono G, Cunningham AA, Fichet-Calvet E, Garry RF, Grant DS, Leach M, et al. A Unified Framework for the Infection Dynamics of Zoonotic Spillover and Spread. PLoS Neglected Tropical Diseases. 2016;10: 1–24. doi:10.1371/journal.pntd.0004957

42. Narat V, Kampo M, Heyer T, Rupp S, Ambata P, Njouom R, et al. Using physical contact heterogeneity and frequency to characterize dynamics of human exposure to non-human primate bodily fluids in central Africa. PLoS Neglected Tropical Diseases. 2018;12: 1–25. doi:10.1371/journal.pntd.0006976

43. Prem K, Cook AR, Jit M. Projecting social contact matrices in 152 countries using contact surveys and demographic data. PLoS Computational Biology. 2017;13: e1005697. doi:https://doi.org/10.1371/journal.pcbi.1005697

44. Prem K, Liu Y, Russell TW, Kucharski AJ, Eggo RM, Davies N, et al. The effect of control strategies to reduce social mixing on outcomes of the COVID-19 epidemic in Wuhan, China: a modelling study. The Lancet Public Health. 2020;5: e261–e270. doi:10.1016/S2468-2667(20)30073-6

45. Craft ME. Infectious disease transmission and contact networks in wildlife and livestock. Philosophical Transactions of the Royal Society B: Biological Sciences. 2015;370. doi:10.1098/rstb.2014.0107

46. Johnson PTJ, de Roode JC, Fenton A. Why infectious disease research needs community ecology. Science. 2015;349. doi:10.1126/science.1259504

47. Boynton PM. Administering, analysing, and reporting your questionnaire. Bmj. 2004;329: 323. doi:10.1136/bmj.329.7461.323-b

48. Berrian AM, Smith MH, van Rooyen J, Martínez-López B, Plank MN, Smith WA, et al. A community-based One Health education program for disease risk mitigation at the human-animal interface. One Health. 2018;5: 9–20. doi:10.1016/j.onehlt.2017.11.002

49. Kalema-Zikusoka G, Kock RA, Macfie EJ. Scabies in free-ranging mountain gorillas (Gorilla beringei beringei) in Bwindi Impenetrable National Park, Uganda. Veterinary Record. 2002;150: 12–15. doi:10.1136/vr.150.1.12

50. Ahebwa WM, van der Duim R, Sandbrook C. Tourism revenue sharing policy at Bwindi Impenetrable National Park, Uganda: A policy arrangements approach. Journal of Sustainable Tourism. 2012;20: 377–394. doi:10.1080/09669582.2011.622768

51. le Polain de Waroux O, Cohuet S, Ndazima D, Kucharski AJ, Juan-Giner A, Flasche S, et al. Characteristics of human encounters and social mixing patterns relevant to infectious diseases spread by close contact: A survey in Southwest Uganda. BMC Infectious Diseases. 2018;18: 1–12. doi:10.1186/s12879-018-3073-1

52. Conservation through public health C. CTPH Annual Report. 2014; 1–16. Available: https://ctph.org/annual-reports/

53. United Nations Educational S and CO. Uganda. 2019 [cited 25 May 2020]. Available: http://uis.unesco.org/country/UG

54. Kamugisha SR, Dobson AE, Stewart AG, Haven N, Mutahunga B, Wilkinson E. A retrospective cross sectional study of the effectiveness of a project in improving infant health in Bwindi, South Western Uganda. Frontiers in Public Health. 2018;6: 1–10. doi:10.3389/fpubh.2018.00290

55. Bishop-Williams KE, Berrang-Ford L, Sargeant JM, Pearl DL, Lwasa S, Namanya DB, et al. Understanding weather and hospital admissions patterns to inform climate change adaptation strategies in the healthcare sector in uganda. International Journal of Environmental Research and Public Health. 2018;15. doi:10.3390/ijerph15112402

56. Sleeman JM, Rooney MB. Determining the Human Diseases Transmissible To the Great Apes of Western Uganda. AAZV and IAAAM Joint Conference. 2000; 36–38.

57. Edwards S, Brooks T. Zoonoses transmissible from non-human primates. 1st ed. OIE Manual of Diagnostic Tests and Vaccines for Terrestrial Animals. 1st ed. OIE; 2018. pp. 1–5. Available: https://www.oie.int/fileadmin/Home/eng/Health_standards/tahm/3.09.11_NONHUMAN_PRIMATES.pdf

58. Homsy J. Ape tourism and human diseases: how close should we get? Tourism. 1999; 70. Available: http://www.igcp.org/library/

59. UWA UWA. The Ultimate Gorilla Experience. 1994 [cited 16 Nov 2020] pp. 1–2. Available: https://www.ugandawildlife.org/explore-our-parks/parks-by-name-a-z/bwindi-impenetrable-national-park

60. Hamilton A, Cunningham A, Byarugaba D, Kayanja F. Conservation in a Region of Political Instability: Bwindi Impenetrable Forest, Uganda. Conservation Biology. 2000;14: 1722–1725. doi:10.1111/j.1523-1739.2000.99452.x

61. Hodd M. East Africa Handbook: The Travel Guide. Footprint Travel Guides. 7th Revise. Footprint Handbooks; 2002.

62. Tumusiime DM, Sjaastad E. Conservation and Development: Justice, Inequality, and Attitudes around Bwindi Impenetrable National Park. Journal of Development Studies. 2014;50: 204–225. doi:10.1080/00220388.2013.841886

63. Wild RG, Mutebi J. Conservation through Community use of Plant Resources: Establishing collaborative management at Bwindi Impenetrable and Mgahinga Gorilla National Parks, Uganda. People and Plants Initiative. 1996; 1–45.

64. Rwego IB. Prevalence of clinical signs in mountain gorillas, Bwindi Impenetrable National Park. 2004.

65. Jena R. Hickey, Eustrate Uzabaho, Moses Akantorana J, Arinaitwe, Ismael Bakebwa, Robert Bitariho, Winnie Eckardt, Kirsten Gilardi, Jacques Katutu, Charles Kayijamahe, Elizabeth M. Kierepka, Benjamin Mugabukomeye, Altor Musema, Henry Mutabaazi, Martha M. Robbins, Benjamin N. Sacks GKZ. Bwindi-Sarambwe 2018 Surveys: monitoring mountain gorillas, other select mammals, and human activities. 2019. Available: https://igcp.org/content/uploads/2020/09/Bwindi-Sarambwe-2018-Final-Report-2019_12_16.pdf

66. Cranfield M, Minnis R. An integrated health approach to the conservation of Mountain gorillas Gorilla beringei beringei. International Zoo Yearbook. 2007;41: 110–121. doi:10.1111/j.1748-1090.2007.00021.x

67. Newman MEJ. Mixing patterns in networks. Physical Review E - Statistical Physics, Plasmas, Fluids, and Related Interdisciplinary Topics. 2003;67: 13. doi:10.1103/PhysRevE.67.026126

68. R Core Team. R: A language and environment for statistical computing. Vienna, Austria: R Foundation for Statistical Computing, Vienna, Austria.; 2020. Available: https://www.r-project.org/

69. Csardi G, Nepusz T. The igraph software package for complex network research. InterJournal. 2006;Complex Sy: 1695.

70. Wickham H. ggplot2 Elegant Graphics for Data Analysis (Use R!). Springer. 2016. doi:10.1007/978-0-387-98141-3

71. Guerrera W, Sleeman JM, Jasper SB, Pace LB, Ichinose TY, Reif JS. Medical survey of the local human population to determine possible health risks to the mountain gorillas of Bwindi impenetrable forest National Park Uganda. International Journal of Primatology. 2003;24: 197–207. doi:10.1023/A:1021410931928

72. Coates J, Swindale a, Bilinsky P. Household Food Insecurity Access Scale (HFIAS) for measurement of food access: indicator guide. Washington, DC: Food and Nutrition Technical …. 2007; Version 3.

73. Rwego IB, Gillespie TR, Isabirye-Basuta G, Goldberg TL. High rates of Escherichia coli transmission between livestock and humans in rural Uganda. Journal of Clinical Microbiology. 2008;46: 3187–3191. doi:10.1128/JCM.00285-08

74. Rwego IB, Isabirye-Basuta G, Gillespie TR, Goldberg TL. Gastrointestinal bacterial transmission among humans, mountain gorillas, and livestock in Bwindi Impenetrable National Park, Uganda. Conservation Biology. 2008;22: 1600–1607. doi:10.1111/j.1523-1739.2008.01018.x

75. Nizeyi JB, Innocent RB, Erume J, Kalema GRNN, Cranfield MR, Graczyk TK. Campylobacteriosis, salmonellosis, and shigellosis in free-ranging human-habituated mountain gorillas of Uganda. Journal of Wildlife Diseases. 2001;37: 239–244. doi:10.7589/0090-3558-37.2.239

76. Graczyk TK, DaSilva AJ, Cranfield MR, Nizeyi JB, Kalema GRNN, Pieniazek NJ. Cryptosporidium parvum Genotype 2 infections, in free-ranging mountain gorillas (Gorilla gorilla beringei) of the Bwindi Impenetrable National Park, Uganda. Parasitology Research. 2001;87: 368–370. doi:10.1007/s004360000337

77. Sleeman A, Jonathan M, Lisa L, James W, Volcans DES. Gastrointestinal Parasites of Mountain Gorillas (Gorilla Gorilla Beringei) in the Parc National Des Volcans, Rwanda. Journal of Zoo and Wildlife Medicine. 2000;31: 322–328. doi:10.1638/1042-7260(2000)031[0322:gpomgg]2.0.co;2

78. Muehlenbein MP, Ancrenaz M. Minimizing pathogen transmission at primate ecotourism destinations: The need for input from travel medicine. Journal of Travel Medicine. 2009;16: 229–232. doi:10.1111/j.1708-8305.2009.00346.x

79. Group TMGVP 2002 EH, MGVP. Risk of Disease Transmission between Conservation Personnel and the Mountain Gorillas: Results from an Employee Health Program in Rwanda. EcoHealth. 2004;1: 351–361. doi:10.1007/s10393-004-0116-4

80. Adams HR, Sleeman JM, Rwego I, New JC. Self-reported medical history survey of humans as a measure of health risk to the chimpanzees (Pan troglodytes schweinfurthii) of Kibale National Park, Uganda. Oryx. 2001;35: 308–312. doi:10.1046/j.1365-3008.2001.00194.x

81. Salerno J, Ross N, Ghai R, Mahero M, Travis DA, Gillespie TR, et al. Human–Wildlife Interactions Predict Febrile Illness in Park Landscapes of Western Uganda. EcoHealth. 2017;14: 675–690. doi:10.1007/s10393-017-1286-1

82. Uganda AIDS Commission. Uganda Population Based HIV Impact Assessment. Uac. 2017; 62–65. Available: https://reliefweb.int/sites/reliefweb.int/files/resources/UPHIA Uganda factsheet.pdf

83. Cleaveland S, Laurenson MK, Taylor LH. Diseases of humans and their domestic mammals: Pathogen characteristics, host range and the risk of emergence. Philosophical Transactions of the Royal Society B: Biological Sciences. 2001;356: 991–999. doi:10.1098/rstb.2001.0889

84. Kalema-Zikusoka G, Rubanga S, Mutahunga B, Sadler R. Prevention of Cryptosporidium and GIARDIA at the human/Gorilla/livestock interface. Frontiers in Public Health. 2018;6: 1–7. doi:10.3389/fpubh.2018.00364

85. Wallis J, Lee DR. Primate conservation: The prevention of disease transmission. International Journal of Primatology. 1999;20: 803–826. doi:10.1023/A:1020879700286

86. Release SDZSP. San Diego Zoo Safari Press Release. San Diego; [cited 28 Jan 2021] p. 2021. Available: https://zoo.sandiegozoo.org/pressroom/news-releases/san-diego-zoo-safari-park-gorillas-recovering-after-sars-cov-2-diagnosis

87. Hope K, Goldsmith ML, Graczyk T. Parasitic health of olive baboons in Bwindi Impenetrable National Park, Uganda. Veterinary Parasitology. 2004;122: 165–170. doi:10.1016/j.vetpar.2004.03.017

88. Chiu CY, Yagi S, Lu X, Yu G, Chen EC, Liu M, et al. A novel adenovirus species associated with an acute respiratory outbreak in a baboon colony and evidence of coincident human infection. mBio. 2013;4: 1–12. doi:10.1128/mBio.00084-13

89. Chen EC, Yagi S, Kelly KR, Mendoza SP, Maninger N, Rosenthal A, et al. Cross-species transmission of a novel adenovirus associated with a fulminant pneumonia outbreak in a new world monkey colony. PLoS Pathogens. 2011;7. doi:10.1371/journal.ppat.1002155

90. (NOHP) TNOHP. Uganda one health strategic plan 2018 - 2022. Ministry of Health (MoH). 2018. Available: http://apps.who.int/medicinedocs/pdf/h2977e/h2977e.pdf

91. Levenson M. Four Poachers Arrested After Killing of Rare Silverback Gorilla in Uganda. 2020: 1. Available: https://www.nytimes.com/2020/06/12/world/africa/rafiki-silverback-gorilla-poachers.html

92. Namara A. From Paternalism to Real Partnership with Local Communities? Experiences from Bwindi Impenetrable National Park (Uganda). Africa Development / Afrique et Développement. 2006;31: 37–66.

93. MacKenzie CA, Hartter J. Demand and proximity: Drivers of illegal forest resource extraction. Oryx. 2013;47: 288–297. doi:10.1017/S0030605312000026

94. Choi BCK, Pak AWP. A catalog of biases in questionnaires. Preventing Chronic Disease. 2005;2: 1–13.

95. Althubaiti A. Information bias in health research: Definition, pitfalls, and adjustment methods. Journal of Multidisciplinary Healthcare. 2016;9: 211–217. doi:10.2147/JMDH.S104807

96. Smith-Greenaway E. Are literacy skills associated with young adults’ health in Africa? Evidence from Malawi. Social Science and Medicine. 2015;127: 124–133. doi:10.1038/jid.2014.371

97. Harrison M. Penetrating the Impenetrable: Establishing profiles and motivations of resource users at Bwindi Impenetrable. 2013; 100.

98. Jones KE, Patel NG, Levy MA, Storeygard A, Balk D, Gittleman JL, et al. Global trends in emerging infectious diseases. Nature. 2008;451: 990–993. doi:10.1038/nature06536

99. Kyampaire. O. The implications of socioeconomic status bordering commnunities on sustainability of natural resources with and adjacent to protected area: the case of Kibale National Park, Uganda. 2004.

100. Berrang-Ford L, Dingle K, Ford JD, Lee C, Lwasa S, Namanya DB, et al. Vulnerability of indigenous health to climate change: A case study of Uganda’s Batwa Pygmies. Social Science and Medicine. 2012;75: 1067–1077. doi:10.1016/j.socscimed.2012.04.016

101. Patterson K, Berrang-Ford L, Lwasa S, Namanya DB, Ford J, Twebaze F, et al. Seasonal variation of food security among the Batwa of Kanungu, Uganda. Public Health Nutrition. 2017;20: 1–11. doi:10.1017/S1368980016002494

102. Twinamatsiko M, Baker J, Harrison M, Shirkhorshidi M, Bitariho R, Wieland M, et al. Equity and poverty alleviation understanding profiles and motivations of resource users and local perceptions of governance at Bwindi Impenetrable National Park. 2014. doi:10.13140/2.1.4853.4409

103. Heesterbeek H, Anderson RM, Andreasen V, Bansal S, DeAngelis D, Dye C, et al. Modeling infectious disease dynamics in the complex landscape of global health. Science. 2015;347. doi:10.1126/science.aaa4339

